# miR-126 modulates neural progenitor cell development

**DOI:** 10.1101/2024.12.27.630551

**Authors:** Jeroen Bastiaans, Michael D. Glendinning, Natalia de Marco Garcia, Heidi Stuhlmann

## Abstract

Noncoding microRNAs (miRNAs) play important roles in controlling signaling pathways by targeting multiple genes and altering their expression levels. MiR-126 is a known endothelial-specific miRNA that regulates vascular integrity and angiogenesis by enhancing proangiogenic actions of VEGF and FGF. MiR-126 is also expressed in several stem cell compartments, including embryonic stem cell (ESC) and hematopoietic stem/progenitor cells (HSPC) where it regulates cell differentiation and quiescence. Here we show that miR-126 is expressed as well in neural progenitor cells where it regulates cell differentiation by targeting genes of the IGF1R signaling pathway including IRS1, PI3K and AKT. Moreover, Hoxa9 and IGF-1 expression by neural progenitor cells is upregulated when miR-126 expression levels are reduced. Our data, implicating a role for miR-126 in manipulating cell development, could open a window of opportunities for clinical purposes and therapeutic strategies to achieve controlling neural progenitor cell behavior.

**Highlights:**

- Knockdown of miR-126 in neural progenitor cells increases cell proliferation
- miR-126 modulates expression levels of HoxA9 and known target genes involved in the IGF1R signaling pathway in neural progenitor cells
- Overexpression of miR-126 results in induced cell differentiation in neural progenitor cells
- miR-126 affects motor neuron development in the thoracic region of the spinal cord
- Knockdown of miR-126 expression results in increased IGF-1 expression

## Introduction

Neural stem and progenitor cells (NSPCs) are multipotent cells in the central nervous system (CNS) that are capable to differentiate into neuronal and glial cells during embryonic development and in the adult brain. NSPC secrete neurotropic factors, growth factors and cytokines that have protective properties for existing neurons (Tang, Yu, and Cheng 2017). These soluble factors direct the progenitor cells to differentiate into specific cells and can be manipulated under in vitro conditions (Okano 2012; Ulrich, Abbracchio, and Burnstock 2012; Kaneko et al. 2000). Controlling the developmental potential of NSPC to give rise to neuronal and glial cells can be a powerful tool for advanced research technologies and treatment strategies for neurological disorders like Parkinson’s Disease, Alzheimer’s Disease, and epilepsy. This may affect millions of patients on a global level (Sugai et al. 2016; Okano 2012; Collaborators et al. 2021; Feigin et al. 2020; Carroll 2019).

Recently, microRNAs (miRNAs) have been recognized as critical epigenetic post-transcriptional regulators involved in neural induction and differentiation (Stappert, Roese-Koerner, and Brustle 2015). miRNAs are non-coding RNA molecules of 20-24 nucleotides in length that reduce gene expression post-transcriptionally by binding to complementary sequences in mRNAs (Bartel 2009; Shukla, Singh, and Barik 2011). miRNAs target about 30-60% of all protein-coding genes in the human genome, are mostly evolutionary conserved and are thought to play an important role in biological processes (Friedman et al. 2009; Griffiths-Jones et al. 2008). Therefore, microRNAs are considered potential candidates for new therapeutic strategies (Alamdari et al. 2021).

In this study, we focus on a novel role for the microRNA miR-126 in neural progenitor cell behavior. miR-126 is highly expressed in endothelial cells where it regulates angiogenic signaling and vascular integrity (Wang et al. 2008; Fish et al. 2008; Kuhnert et al. 2008). Mechanistically, miR-126 enhances the proangiogenic actions of vascular endothelial growth factor (VEGF) and fibroblast growth factor (FGF) through repression of Sprouty Related EVH1 Domain Containing (SPRED)-1 and Phosphoinositide-3-Kinase Regulatory Subunit 2 (PI3KR2/p85-beta) (Wang et al. 2008; Huang et al. 2020; Kuhnert et al. 2008). MiR-126 is located in intron 7 of its host gene *Egfl7,* which encodes an extracellular matrix-associated and secreted angiogenic factor (Fitch et al. 2004; Campagnolo et al. 2005; Parker et al. 2004; Soncin et al. 2003; Nichol and Stuhlmann 2012). *Egfl7* and miR-126 share upstream regulatory elements for their expression in the endothelial cell lineage (Wang et al. 2008; Bambino et al. 2014).

In addition to endothelial cells, miR-126 is expressed by other cell types including trophoblasts, hepatocytes, and in several stem cell compartments including embryonic stem cells (ESC), trophoblast stem cells (TSC) and hematopoietic stem/progenitor cells (HSPC) (Lechman et al. 2012; Sharma et al. 2019; Ryu et al. 2011). miR-126 plays an important role in HSPC self-renewal and quiescence through targeting multiple genes within the PI3K signaling pathway (Lechman et al. 2012). Several reports show that miR-126 modulates expression of key components of the Insulin Like Growth Factor 1 (IGF-1) and PI3K signaling pathway in various cell types (Qu et al. 2019; Lu et al. 2018; Lechman et al. 2012; Rouigari et al. 2018; Sharma et al. 2019; Ryu et al. 2011).

In this study, we investigated the role of miR-126 in the NSPC lineage, specifically its function in NPC differentiation and in HoxA9 signaling. While its host gene *Egfl7* is expressed by NSCs and adult neurons and has been implicated in neurogenesis (Bicker et al. 2017), it was not known if miR-126 is expressed in NPCs and what potential role it plays during neurogenesis.

## Results

### miR-126 affects proliferation of human iPSC-derived NPCs

Human iPSC-derived neural progenitor cells (NPCs) were provided by Dr. Shuibing Chen (Weill Cornell Medicine) and maintained in NPC maintenance medium for no longer than 8 cell passages. We compared levels of miR-126 expression in human NPCs to its expression in human umbilical vein endothelial cells (HUVECs) and human skin fibroblasts by qRT-PCR. Relative levels of miR-126 were about 100-fold higher in NPCs compared to fibroblasts and 100-fold lower compared to HUVECs (Figure 1A). To investigate the role of miR-126 in NPCs, we performed overexpression and knock-down experiments. Lentiviral constructs to overexpress (miR-126/OE) or knock-down miR-126 (miR-126/KD), and their respective control constructs (control/OE and control/KD), were provided by John E. Dick (University of Toronto, Canada) (Lechman et al. 2012). The lentiviral construct for overexpression of miR-126 and its control (miR-126/OE; control/OE) contain a RFP marker, and the lentiviral construct for knock-down miR-126 and its control (miR-126/KD; control/KD) contain a GFP marker (Lechman et al. 2012). Transduction of NPC with miR-129/OE lentivirus increased miR-126 expression almost 50-fold (*P* = 0.02), while miR-126/KD lentivirus almost completely depleted miR-126 expression (*P* = 0.006) (Figure 1B), with approximately 80% of the cells infected as judged by their expression of RFP or GFP. Co-staining of the transduced NPCs for the proliferation marker Ki67 and RFP or GFP showed an about 2-fold increase in proliferation in miR-126/KD transduced cells compared to the control transduced cells, whereas no difference in proliferation was detected between miR-126/OE and control/OE transduced NPC (Figure 1C, D). Co-staining for the apoptotic marker Cleaved Caspase-3 and RFP or GFP did not reveal any changes in cell death in cells with overexpression or knockdown of miR-126 (Supplemental Figure 1).

**Figure 1:**
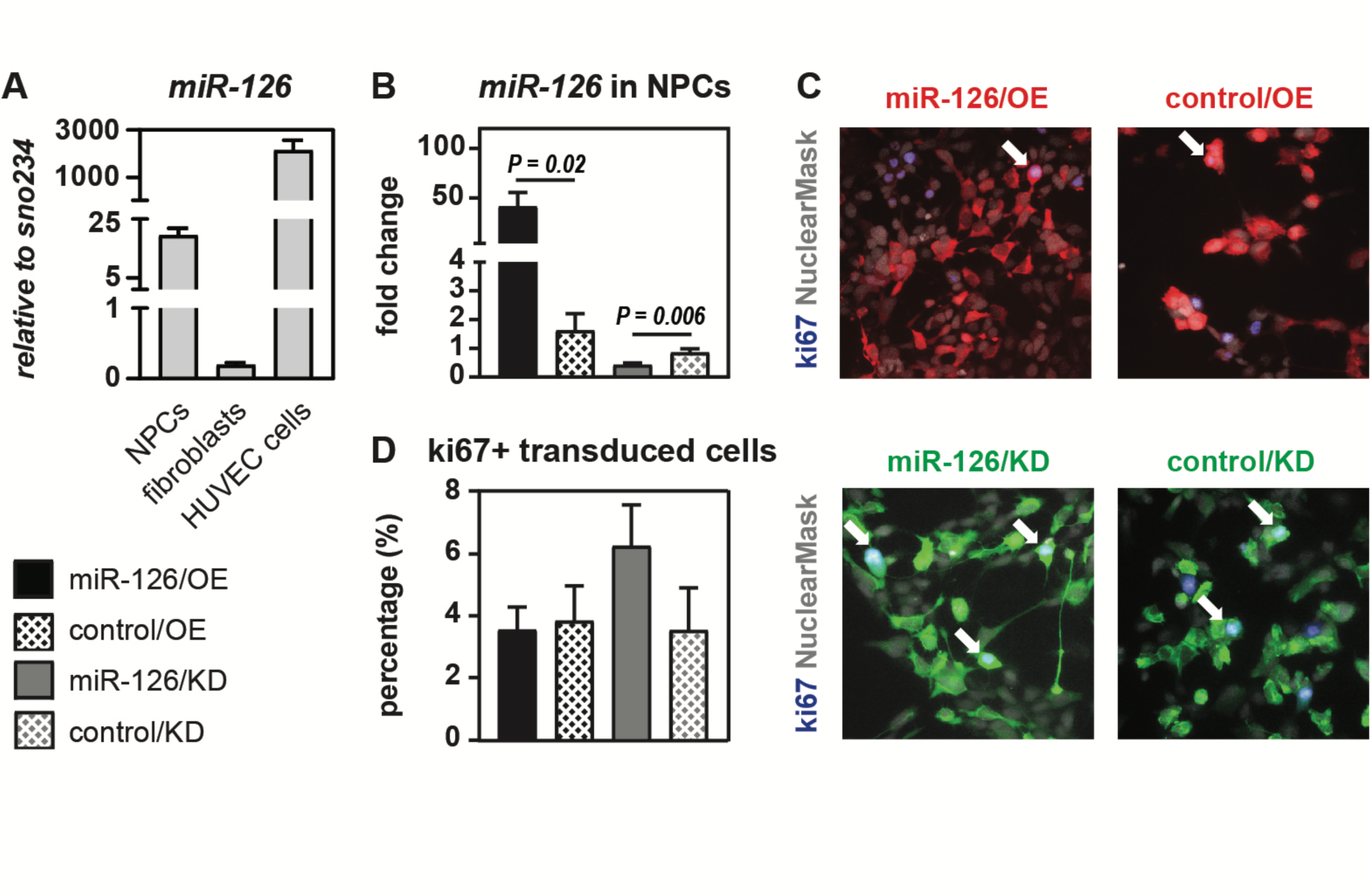
miR-126 is expressed in NPCs and knock-down of its expression results in increased cell proliferation. **A.** Expression levels of miR-126 in NPCs, human skin fibroblasts, and HUVECs by RT-PCR. Levels of expression are shown relative to expression of sno234. **B.** Expression of miR-126 in NPCs infected with lentivirus to overexpress (miR-126/OE), its empty vector control (control/OE), and lentivirus to knock-down miR-126 (miR-126/KD), and its empty vector control (control/KD), respectively. MiR-126 expression levels were determined by qRT-PCR (n = 8) and normalized to uninfected NPC controls. miR-126/OE transduced NPCs showed significantly elevated levels of miR-126 (*P* = 0.02), while miR-126/KD transduced NPCs showed significantly reduced levels of miR-126 (*P* = 0.006). **C.** Representative images of NPC infected with miR-126/OE or control/OE (top row), and NPC infected with miR-126/KD or control (bottom row). Proliferating cells were identified by Ki67 staining, and proliferating transduced cells were discriminated from non-transduced cells by double staining for RFP (top row) or GFP (bottom row). Images were taken at 20x magnification. **D.** Quantification of miR-126 controlled proliferation of transduced NPCs by counting cells positive for Ki67 and RFP, or Ki67 and GFP, respectively (n=4).

### miR-126 modulates expression of *HOXA9* and regulators in the IGF1R signaling pathway in NPCs

Previous studies reported that miR-126 targets genes in the IGF1R signaling pathway in endothelial cells (ECs) and hematopoietic stem/progenitor cells (HSPCs) (Lechman et al. 2012; Lu et al. 2018; Wang et al. 2008; Wu et al. 2016). Using lentivirus vectors for knockdown of miR-126, we demonstrate that RNA expression levels of several miR-126 target genes that are components of IGFR1 signaling were significantly upregulated in NPCs, including *IRS1*, *AKT* and *HOXA9* (Figure 2A). Lentiviral overexpression of miR-126 in NPCs had no effect on the expression levels of these genes (Figure 2A). Interestingly, the HOXA9 gene is transcribed into two mRNAs isoforms. The miR-126 target sequence is located in the second exon, which encodes the homeodomain, of isoform 1 of the HOXA9 transcript rather than in its 3’ UTR (Shen et al. 2008). An alternatively spliced isoform 2 of HOXA9, termed splice variant 2, lacks the second exon that contains the homeodomain and the miR-126 target sequence (Stadler et al. 2014). Our results show that NPCs exclusively express splice variant 1 of HOXA9 that can be targeted by miR-126 (Supplemental Figure 2).

**Figure 2:**
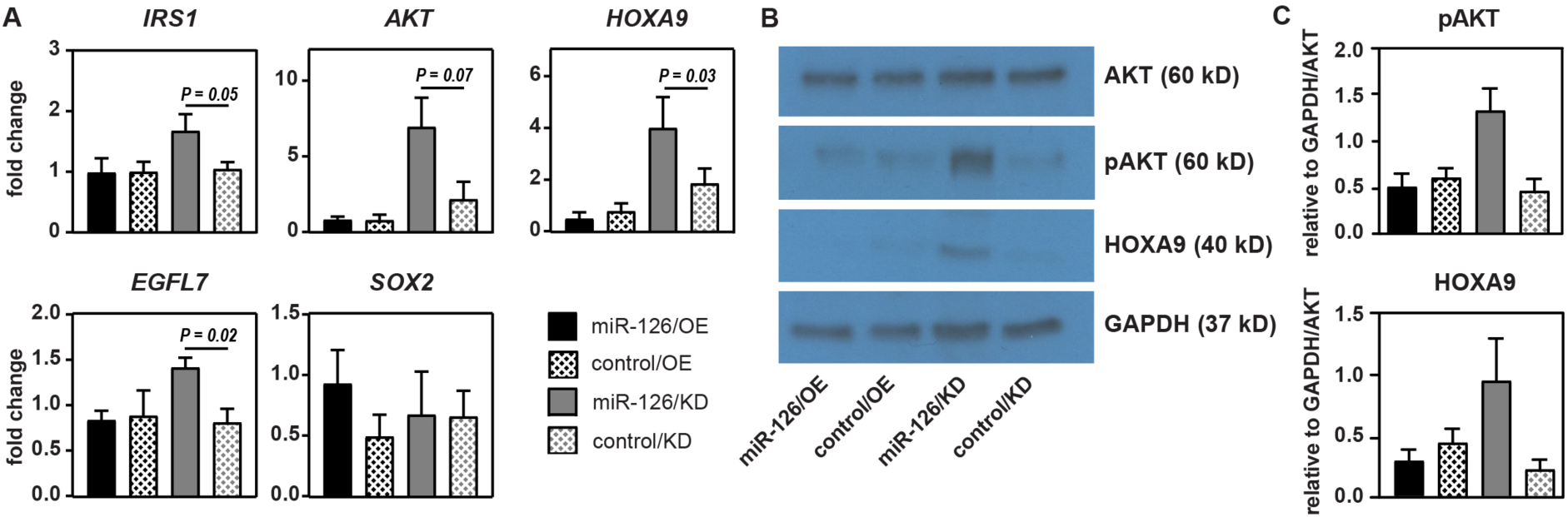
miR-126 targets genes in the IGF1R-signaling pathway in NPCs. **A.** NPCs were transduced with lentivirus to overexpress (miR-126/OE) or knock-down (miR-126/KD) miR-126, and their respective controls (control/OE and control/KD). Expression levels of *IRS1, AKT, HOXA9, EGFL7* and *SOX2* mRNA were determined by qRT-PCR (n = 4) and normalized to levels in uninfected NPCs. Knock-down of miR-126 in NPCs resulted in significantly increased expression of *IRS1* (P = 0.05), *AKT* (P = 0.07), *HOXA9* (P = 0.03) and to a lesser extent of *EGFL7* (P = 0.02) compared to its control, whereas no changes were observed upon overexpression of miR-126. *SOX2* mRNA expression levels were not affected by the overexpression or knock down of miR-126. **B.** Western blot analysis to shows levels of protein expression for AKT (60 kD), pAKT (60 kD), HOXA9 (40 kD) and GAPDH (37 kD) in transduced NPCs. **C.** Quantification of band intensities on western blots show that knock-down of miR-126 in NPCs results in increased levels of pAKT and HOXA9 protein (n=4).

The host gene of miR-126, *EGFL7*, has been reported to be a potential target gene for miR-126 (Sun et al. 2010). Lentiviral knock-down of miR-126 in human NPCs resulted in a small but significant increase of EGFL7 expression (Figure 2A). However, the *EGFL7* 3’ UTR does not contain a miR-126 target sequence, and it remains unknown whether the effect of miR-126 on EGFL7 expression is a direct or an indirect one. miR-126 has been reported to inhibit SOX2 expression in multiple gastric cancer cell lines and in mouse embryonic stem cells (Otsubo et al. 2011). However, no change of *SOX2* expression was observed in NPCs transduced with miR-126/KD lentivirus (Figure 2A).

Next, we investigated protein expression levels of AKT, pAKT and HOXA9 in NPCs that were transduced with mir-126/OE, miR-126/KD and control lentiviruses. Western blot analysis showed pAKT and HOXA9 bands at increased intensity in NPCs with miR-126 knock-down (Figure 2B). Quantification of pAKT and HOXA9 band intensities on western blots and normalization against AKT and GAPDH showed increased levels of pAKT and HOXA9 protein (Figure 2C), consistent with our finding of increased AKT and HOXA9 mRNA expression (Figure 2A). Together, our results indicate that miR-126 inhibits the expression of key regulators of the IGF1R pathway in NPCs, including IRS1, AKT and HOXA9.

### miR-126 affects human motor neuron development in vitro

Members of the HOX gene family function as important transcriptional regulators of embryonic development. HOXA9 is expressed by a subset of motor neurons in the thoracic region of the spinal cord (Dasen, Liu, and Jessell 2003; Philippidou and Dasen 2013) To investigate if miR-126-mediated changes of HOXA9 expression levels affect motor neuron development, we infected NPCs with miR-126/OE or miR-126/KD lentivirus, and their respective controls, and induced their differentiation into motor neurons by culture for 3 days in motor neuron differentiation medium. Differentiating NPCs are expected to show a decrease in expression of progenitor cell markers and an increase in motor neuron specific markers. To account for heterogeneity of the NPCs in culture and for the short time allowed for their differentiation, PAX6 and OLIG2 were selected as NPC markers, NKX6.1 which is expressed in motor neuron progenitors and retained by subsets of postmitotic motor neurons and ISL1 is a pan-motor neuronal marker (Figure 3A) (Davis-Dusenbery et al. 2014; De Marco Garcia and Jessell 2008). First, the extent of differentiation was determined by analyzing RNA expression levels by qRT-PCR in undifferentiated and differentiated NPCs in the absence of lentivirus transduction (Figure 3B). After 3 days in differentiation medium, no changes in *PAX6* or *OLIG2* expression levels were detected in the non-transduced NPCs, indicating that the majority of cells in culture remain as undifferentiated NPCs. However, *NKX6.1* and *ISL1* expression levels were significantly induced, indicating the presence of cells that differentiated along the NPC - motor neuron axis (Figure 3B).

**Figure 3:**
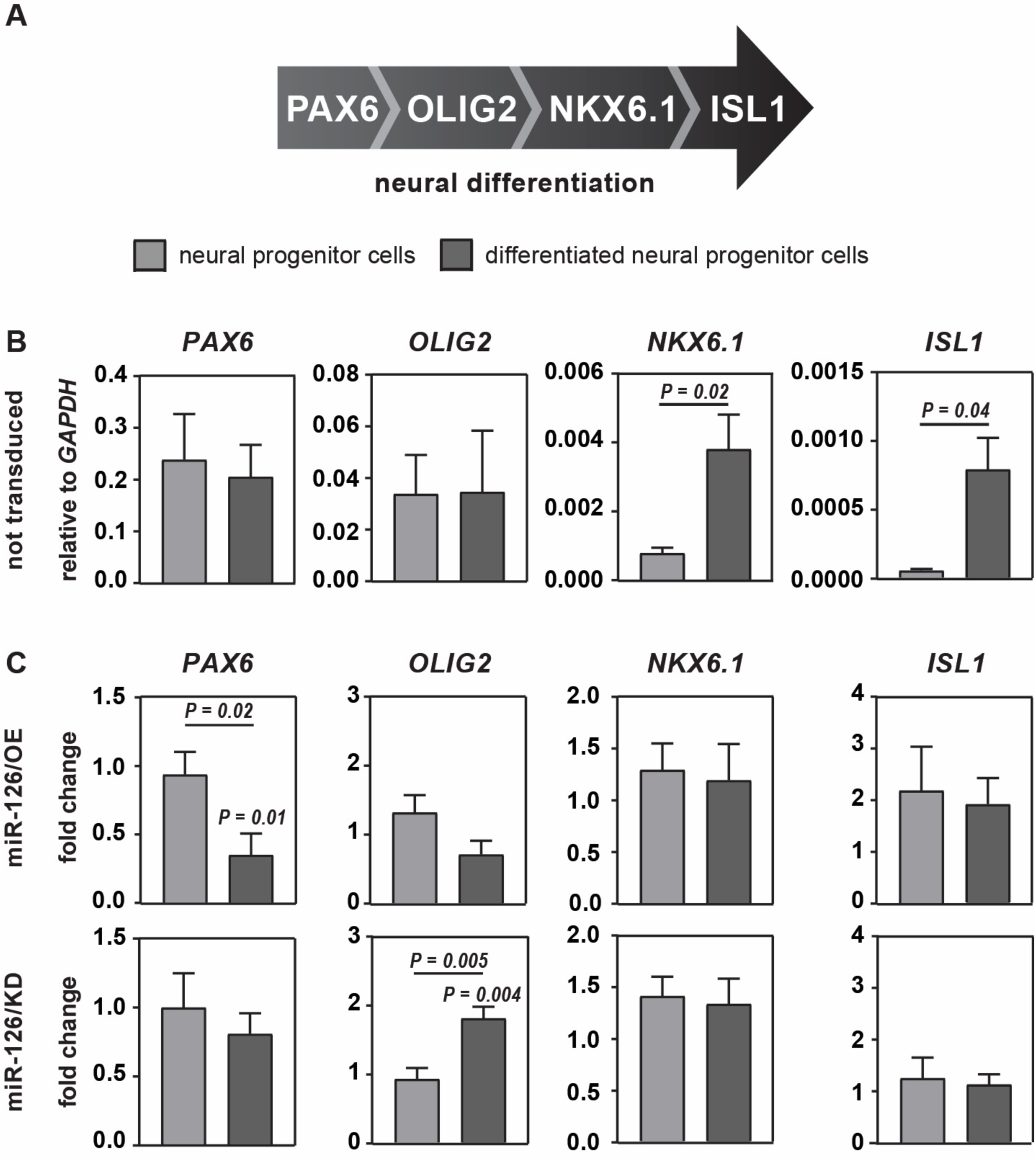
miR-126 affects neurogenesis in human NPCs. **A.** Schematic overview of neural marker expression during differentiation of neural progenitor cells. **B.** Expression levels of neural progenitor and motor neuron markers *PAX6*, *OLIG2*, *NKX6.1* and *ISL1* in undifferentiated NPCs and in cells after 3 days in motor neuron differentiation medium were determined in the absence of lentivirus transduction by qRT-PCR (n = 6). **C.** Expression levels of neural progenitor and motor neuron markers *PAX6*, *OLIG2*, *NKX6.1* and *ISL1* in undifferentiated NPCs transduced with lentivirus to overexpress (miR-126/OE) and knock-down (miR-126/KD) miR-126, and controls (control/OE and control/KD), respectively, and in cells after 3 days of culture in motor neuron differentiation medium. All expression levels were calculated relative to expression of GAPDH. Expression levels are shown as fold-change in differentiating transduced NPCs compared to the undifferentiated transduced NPCs.

To determine how miR-126 affects this process, NPCs were transduced with miR-126/OE or miR-126/KD, and their respective controls. Subsequently, transduced NPCs were cultured for three days in parallel in NPC maintenance medium and in motor neuron differentiation medium. *PAX6*, *OLIG2*, *NKX6.1* and *ISL1* mRNA expression levels in miR-126/OE or miR-126/KD transduced cells were normalized to their respective empty vector controls and compared between differentiated and undifferentiated NPCs (Figure 3C). Here, overexpression of miR-126 in differentiated NPCs resulted in reduced levels of *PAX6* expression compared to undifferentiated NPCs (*P* = 0.02) and their control (*P* = 0.01). A similar change, albeit not reaching significance, was seen for *OLIG2* expression in differentiated NPCs overexpressing miR-126 (Figure 3C), indicating that overexpression of miR-126 reduces expression of early progenitor markers and pushes NPCs towards differentiation. Conversely, knock-down of miR-126 resulted in upregulated *OLIG2* expression in differentiated NPCs compared to undifferentiated NPCs (*P* = 0.005) and their controls (*P* = 0.004), whereas *PAX6*, *NKX6.1* and *ISL1* expression levels remained unchanged (Figure 3C). While PAX6 is considered a common neural stem cell marker, *OLIG2* is the first marker to be expressed by neurons committed to the motor neuron fate (Sun et al. 2011; Thakurela et al. 2016). *OLIG2* also promotes neural stem cell and progenitor cell proliferation (Ligon et al. 2007). Together, these results suggest that overexpression of miR-126 decreases proliferation and promotes differentiation of NPCs, whereas knockdown miR-126 promotes proliferation and blocks NPC differentiation to the motor neuron fate.

### miR-126 affects motor neuron development in the murine spinal cord

Next, we investigated how miR-126 affects motor on neuronal development *in vivo*, using a mouse model with a global, targeted deletion of miR-126 (Wang et al. 2008; Sharma et al. 2019). Specifically, to examine how miR-126 loss-of-function affects differentiation of NPCs in the thoracic region of the spinal cord that expresses Hoxa9, sagittal sections from E13.5 wild type, miR-126^+/-^ and miR-126^-/-^ embryos were stained by immunofluorescence (IF) for PAX6, OLIG2, ISLET1, SOX2 and HOXA9 (Figure 4A). Sections of the forebrain and ganglionic eminence from the same mice were used as controls since they express PAX6 and OLIG2 but do not express HOXA9 at this developmental stage (Figure 4B). To exclude possible effects resulting from a vascular mutant phenotype in the experimental embryos, sections were immunostained for the endothelial marker CD31 (Figures 4A and 4B). Representative sections (n = 3 for each genotype) are shown in Figure 4A and B, and quantification of IF signal intensities are shown in Figure 4C and D. Strikingly, a shift of HOXA9 positive cells from the ventral spinal cord (left side of image) to the dorsal spinal cord (right side of image) was observed in miR-126^-/-^ embryos, and to a lesser extent in miR-126^+/-^ embryos (Figure 4A), suggesting an expansion and migration of spinal cord motor neurons. Quantification of IF intensities showed increased expression of PAX6, OLIG2 and HOXA9 in the spinal cords of miR-126^-/-^ embryos as compared to wild type embryos (Figure 4C). These results are consistent with our findings in human iPSC-derived human NPCs (Figures 2 and 3). Furthermore, Islet1 expression is reduced in the spinal cords of miR-126^-/-^ embryos compared to wild type embryos, and expression of Sox2 is unchanged (Figure 4C), consistent with our results for *SOX2* RNA expression in the transduced NPCs (Figure 2). In contrast, expression patterns of PAX6 and OLIG2 in brains from wild type, miR-126^+/-^ and miR-126^-/-^ mice look similar and no differences in their IF signal intensity were detected (Figure 4B, D), suggesting that *Pax6* and *Olig2* are not under direct control of miR-126. No changes in vascular endothelial CD31 expression were detected between wild-type and mutant embryos (Figure 4A-D). Together, our results suggest that Hoxa9 is a relevant direct target of miR-126 and that miR-126-induced changes in Hoxa9 expression levels and its localization in the spinal cord may be critical for motor neuron development *in vivo*.

**Figure 4:**
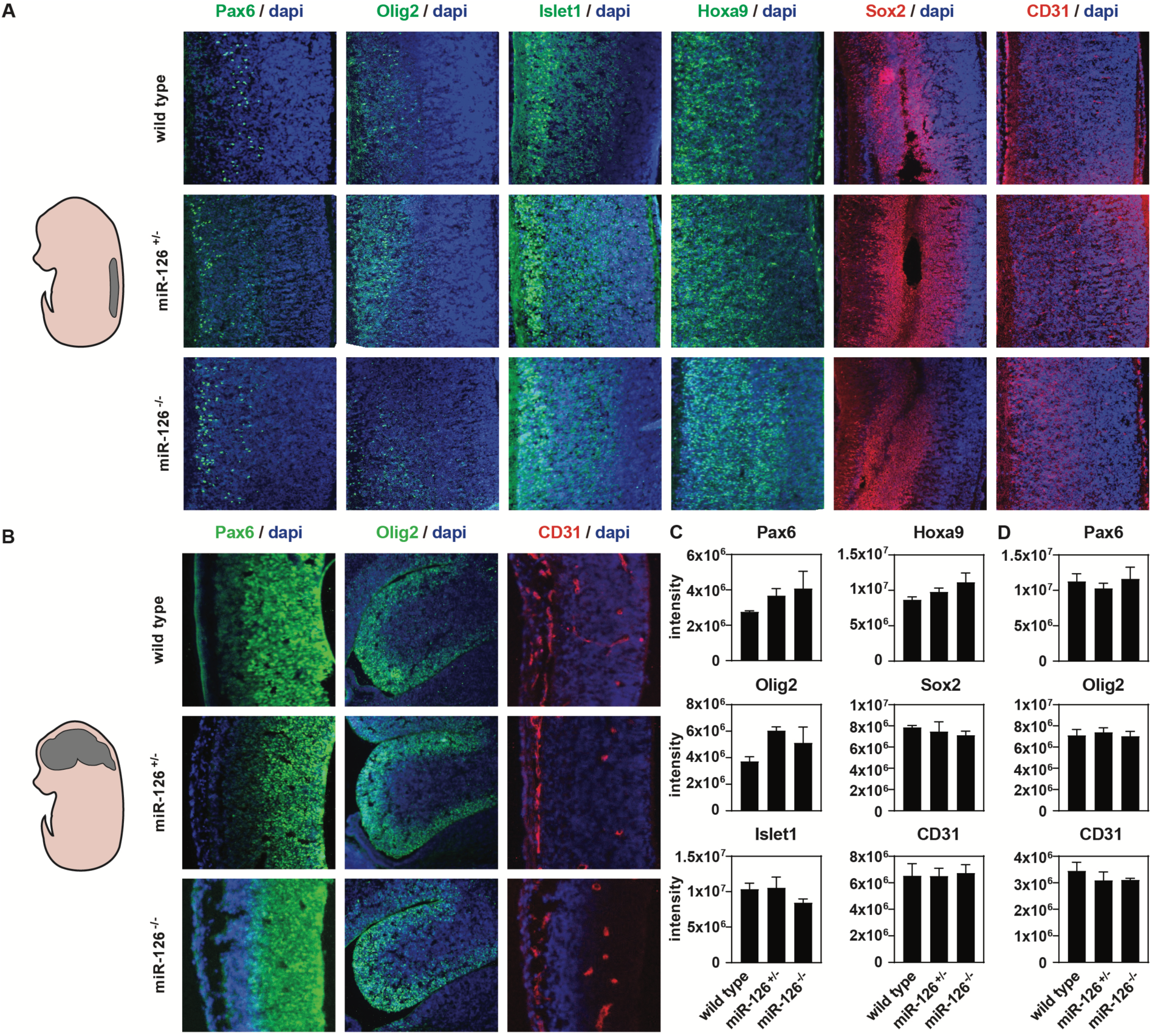
HoxA9 is a relevant target for miR-126 in the developing murine spinal cord. **A,B.** Sagittal tissue sections of the thoracic region of the spinal cord **(A)** and forebrain **(B)** from E13.5 wild type, miR-126^+/-^ and miR-126^-/-^ embryos were immunostained using Pax6, Olig2, Islet1, Hoxa9, Sox2 and CD31 antibodies. Sections for each genotype were matched as best as possible, with images shown being representative for n=3 embryos per genotype. **C**,**D.** Quantification of intensity of fluorescence signals for Pax6, Olig2, Islet1, HoxA9, Sox2 and CD31 in spinal cords **(C)** and forebrains **(D)**, using ImageJ.

### miR-126 knockdown affects IGF1 expression by NPCs

In certain types of leukemia, increased levels of HOXA9 induce an increase of IGF1 expression and of its receptor IGFR1R (Steger et al. 2015; Whelan, Ludwig, and Bertrand 2008). IGF1 is a secreted signaling factor of the PI3K pathway and plays critical roles during embryogenesis (Lewitt and Boyd 2019; Teng, Jeng, and Shyu 2018). Here, we examined *IGF1R* and *IGF1* mRNA expression in NPCs transduced with lentivirus vectors. *IGF1* mRNA expression levels were increased in miR-126/KD transduced NPCs when compared to its control, whereas *IGF1R* expression levels were not affected. No changes in *IGF1* or *IGF1R* expression were observed in NPCs transduced with miR-126/OE (Figure 5A). Finally, we measured IGF1 protein present in culture supernatants of transduced NPCs 24 hours after of passaging the cells. To detect free IGF1 that remains unbound to Insulin-like growth factor-binding proteins (IGFBPs) present in the supernatant, we pre-saturated the supernatants with recombinant human IGF2 (Baxter 2001) prior to performing an ELISA assay. Secreted IGF1 was detected at significantly increased levels in the culture supernatant of miR-126/KD NPCs, whereas no changes were observed for miR-126/OE NPCs (Figure 5B). Together, our results indicate that knock-down of miR-126 in NPC leads to an increase in HOXA9 expression and results in increased levels of secreted IGF1 protein (Figure 6).

**Figure 5:**
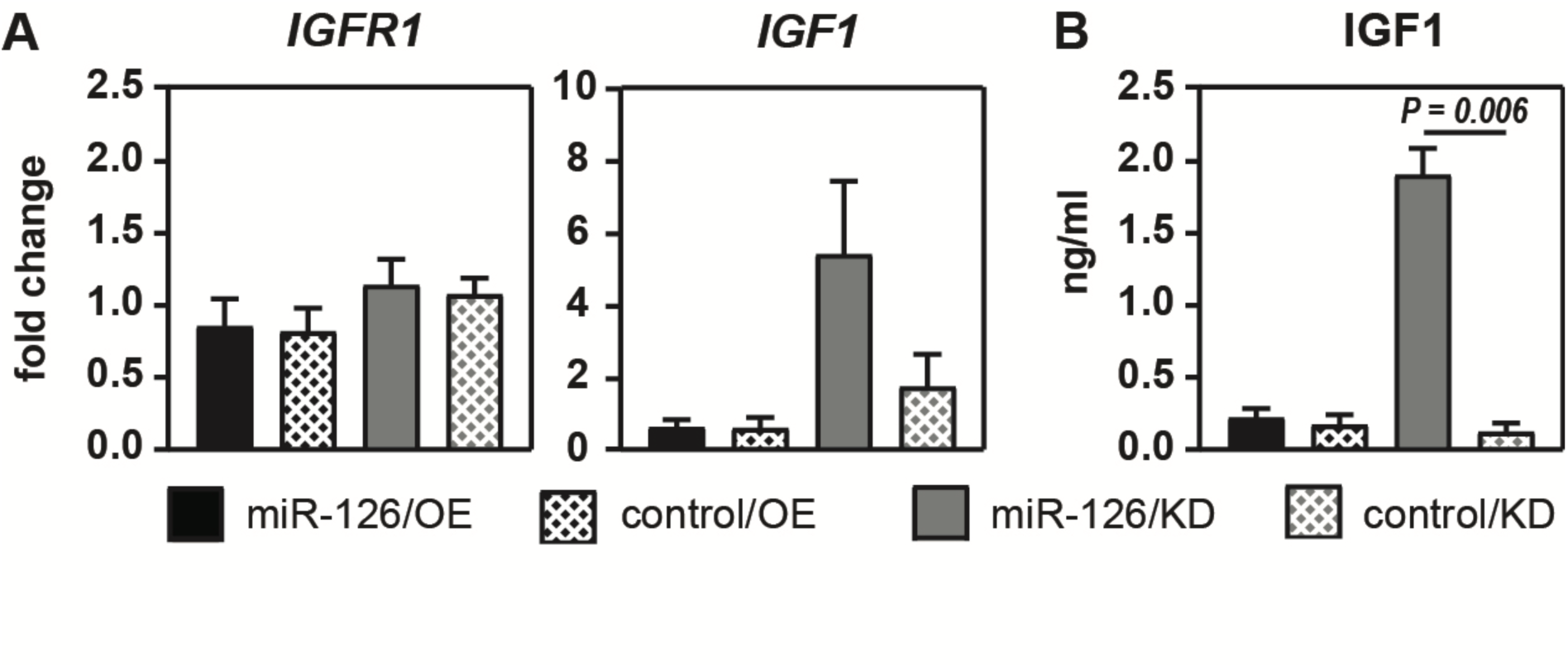
Knockdown of miR-126 in NPCs induces increased levels of IGF-1 expression. **A,B.** NPCs were transduced with lentivirus to overexpress (miR-126/OE), or knock-down (miR-126/KD) miR-126, and the respective controls (control/OE and control/KD). **(A)** *IGFR1* and *IGF-1* mRNA expression levels were determined by qRT-PCR (n = 4). **(B)** IGF1 protein levels from culture supernatants that were collected 24 hours after lentivirus transduction and pre-saturated with recombinant human IGF-2 were determined by ELISA assay.

**Figure 6:**
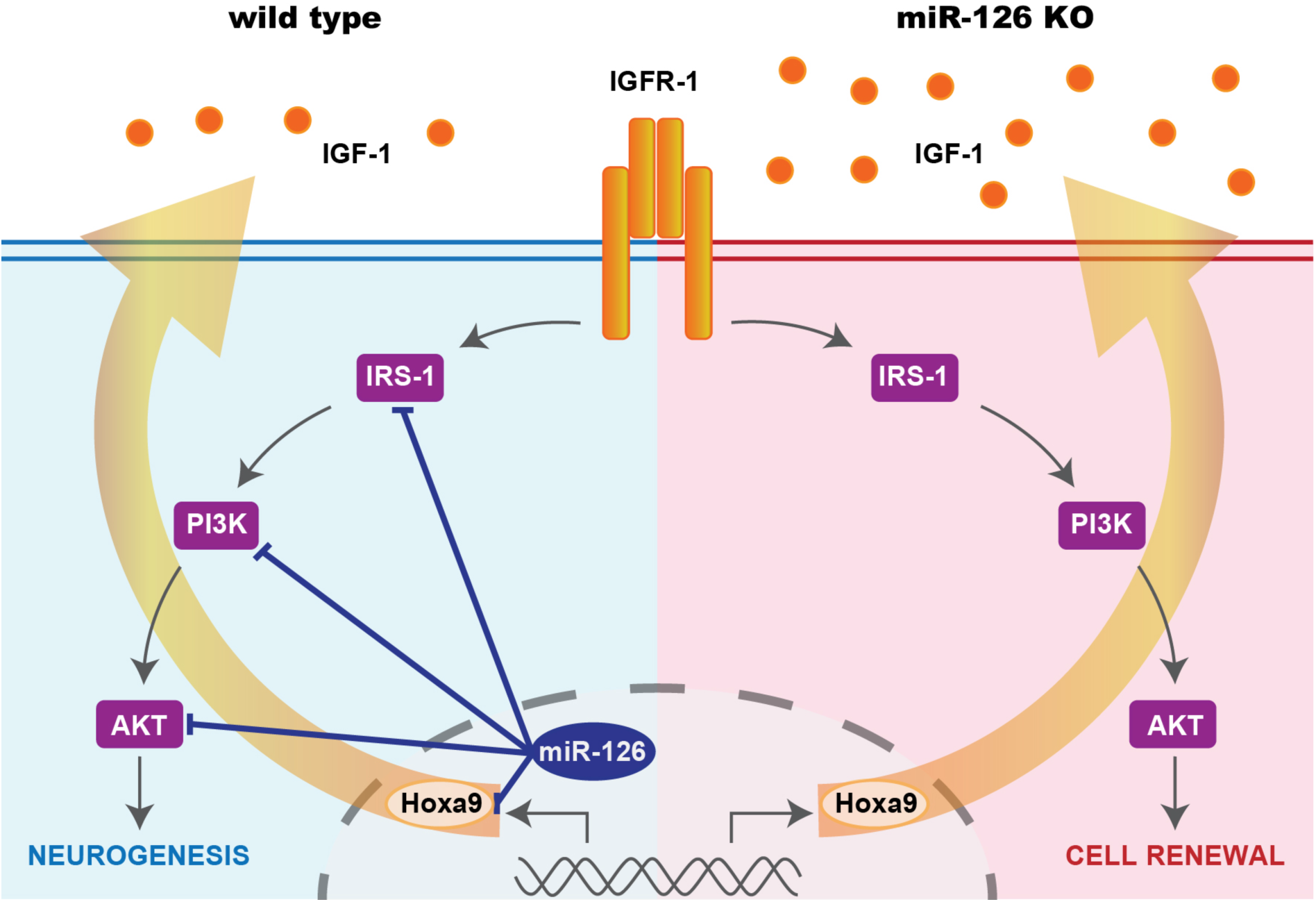
Proposed model. Our model proposes that miR-126 in NPCs contributes to both neurogenesis and cell renewal by modulating the IGF1R pathway. An increase of miR-126 shifts the cells towards neurogenesis, while a decrease of miR-126 results in NPC proliferation/renewal. We hypothesize that in developing motor neurons, loss of miR-126 indirectly results in increased levels of IGF-1 through an increase of Hoxa9 expression, which contributes to the autocrine loop in IGF1R signaling and cell renewal.

## Discussion

iPSC-derived NPCs offer a wide range of opportunities to study developing neurons and serve as models to study neurological and neurodegenerative diseases. In this study we focused on the role of miR-126 in NPC development and its effect on *HOXA9*, a known target for miR-126 (Shen et al. 2008) and genes in the IRS-1/PI3K pathway. miR-126 was first decribed as an endothelial-specific microRNA with a crucial role in developmental angiogenesis and lymphangiogenesis in mice and zebrafish through its repression of SPRED-1 and PI3KR2 (Wang et al. 2008; Huang et al. 2020; Fish et al. 2008; Kontarakis et al. 2018). More recently, miR-126 expression was reported in several types of stem and progenitor cells (Lechman et al. 2012; Sharma et al. 2019; Ryu et al. 2011) including HSPC, where miR-126 sets a threshold for HSC activation through controlling multiple targets of PI3K signaling pathway (Lechman et al. 2012). This prompted us to investigate whether miR-126 is also expressed by NPCs and if it would affect their proliferation and differentiation by targeting genes in the PI3K pathway. Knowledge of how to control this mechanism would be favorable for novel treatment strategies and therapeutical approaches in neurological disorders.

Results we present in this study uncover a novel role for miR-126 in neurogenesis. miR-126 is expressed in human NPCs, and it affects neural cell development by regulating several genes in the IRS1 / PI3K pathway and by directly targeting HOXA9. Lentivirus-mediated overexpression of miR-126 skewed the NPCs towards cell differentiation, whereas knockdown of miR-126 promoted NPC proliferation. Reducing miR-126 in NPCs leads to increased expression of IRS1, AKT and HOXA9, and loss-of miR-126 in mouse embryos results in expanded areas of HOXA9 expressing spinal cord neurons. To our knowledge, this is the first study reporting on the molecular pathways regulated by miR-126 in neural development and how this affects HOXA9 expressing neurons. Previous reports showed that exosomes containing miR-126 promote neurogenesis and angiogenesis and reduce apoptosis in rats after spinal injury (Huang et al. 2020), and that VEGF expression in retinal neural stem cells is regulated by miR-126 by targeting SPRED-1 (Ye et al. 2018).

MicroRNAs function as post-transcriptional regulators and often have moderate effects on the expression of their target genes. Consistent with these findings, some of our experiments using miR-126 overexpression or knockdown in NPCs, and miR-126 knockout in murine spinal cord, show a clear trend in gene expression changes without reaching significance when compared to their controls.

HOXA9 expression was significantly increased in NSCs transduced with miR-126 knockdown lentivirus. In miR-126 knockout mice, increased HoxA9 expression and a shift of HOXA9 positive neurons towards the dorsal side of the spinal cord was detected. Hoxa9 mRNA is a direct target of miR-126. The miR-126 target sequence is located within the Hox domain in exon 2 of the full gene’s transcript and is absent in a truncated version of an alternative spliced transcript (Stadler et al. 2014). We show that NPCs express the full-length transcript of Hoxa9 mRNA. Hoxa9 belongs to the family of Hox genes that control the body plan of embryonic development along the head-tail axis. Hox genes encode transcription factors that contain conserved DNA sequences known as the Hox domain and promote cell division, cell adhesion, apoptosis, and cell migration (Nolte, De Kumar, and Krumlauf 2019; Sanchez-Herrero 2013). Hoxa9 is expressed by stem cells and developing neurons in the thoracic region of the spinal cord (Dasen, Liu, and Jessell 2003; Guthrie 2004; Bhatlekar, Fields, and Boman 2018) where it is essential for the development of local motor neurons. Several studies suggest that that Hoxa9 paralogs, including Hoxb9, Hoxc9 and Hoxd9, can compensate for loss-of Hoxa9 (Jung et al. 2010) (Chen and Capecchi 1997, 1999). It is possible that the lack of an overt motor neuron phenotype in miR-126 KO mice is due to compensatory effects of Hox paralogs. More recent data showed that misexpression of Hoxa9 results in the conversion of limb innervating motor neurons in the lateral motor column to sympathetic ganglia innervating motor neurons in the preganglionic column and that this can repress the expression of Hox4-8 genes in the brachial region (Jung et al. 2010; Dasen, Liu, and Jessell 2003). Therefore, it is possible that increased and mis-localized Hoxa9 expression in the spinal cords of miR-126 KO mice could affect the development of limb innervating motor neurons and preganglionic column motor neurons. Understanding of the consequences of altered Hoxa9 expression pattern in the spinal cord on the development of motor neurons, and on motor skills and behavior in adults, remains to be examined in detail.

Hoxa9 has been described to have a poor prognosis in acute leukemia (Shen et al. 2008; Collins and Hess 2016; Bhatlekar, Fields, and Boman 2018). Elevated levels of miR-126 reduced the expression of Hoxa9 through binding to a region in the Hox domain of its transcript in immortalized bone marrow cells, while inhibition of miR-126 increased expression levels of Hoxa9 in F9 cells (Shen et al. 2008). Furthermore, in B-lineage Acute Lymphoblastic Leukemia (ALL), IGF1-R expression is induced via Hoxa9 and IGF1, a key component in hematopoietic transformation, was shown to be a direct target of Hoxa9 (Whelan, Ludwig, and Bertrand 2008; Steger et al. 2015). The increased levels of IGF-1 protein detected in culture supernatants from miR-126/KD transduced NPCs suggest that IGF-1 expression in NPCs is under control of Hoxa9 as well.

MiR-126 is also expressed in pathologies of fully developed, mature neurons of patients with Schizophrenia, Parkinson’s, and Alzheimer’s Disease (Kim et al. 2014; Kim et al. 2016; Sonntag, Woo, and Krichevsky 2012). Because altered growth factor signaling has been associated with age-related neuronal dysfunction and neurodegenerative diseases like Parkinson’s and Alzheimer’s disease, and since miR-126 regulates key components of IGF1 signaling, miR-126 is likely to contribute to neural pathologies (Castilla-Cortazar et al. 2020; Lewitt and Boyd 2019; Yang et al. 2018; Castelli et al. 2019; Lechman et al. 2012; Sharma et al. 2019). In fact, increased expression of miR-126 was found in dopaminergic neurons of patients with Parkinson’s disease where it has a neurotoxic affect by impairing PI3K, AKT and ERK signaling, while inhibition of miR-126 shows a neuroprotective effect (Kim et al. 2014). Similarly, in a mouse model for familial Alzheimer’s disease, increased levels of miR-126 and Abeta1-42 affected regulators of the IGF1 signaling pathway and resulted in neurotoxic effects (Kim et al. 2016). Therefore, miR-126 is a potential candidate for therapeutical applications. Our model proposes a role for miR-126 in balancing neural development and cell proliferation by regulating genes in the PI3K pathway. Through targeting Hoxa9, miR-126 may control IGF-1 expression, closing the loop by regulating to PI3K pathway signaling in a IGF1R-dependent manner (Figure 6).

## Author contribution

Conceptualization and methodology: J.B., N.dM.G. and H. S.; investigation: J.B. and M.D.G.; writing – original draft: J.B. and H.S.; writing – review and editing: J.B., N.dM.G. and H.S.

## Acknowledgements

The authors would like to thank Drs. Shuibing Chen and Alice Giani (Weill Cornell Medicine) for providing us with iPSC-derived neural progenitor cells, Dr. John E. Dick (University of Toronto, Canada) for providing us with the lentiviral constructs for overexpression and knock-down of miR-126, Drs. Nina Pipalia and Linda Sasset (Weill Cornell Medicine) for fibroblast and HUVEC RNAs, and Takumi Otsuka for technical support.

## Experimental Procedures

### miR-126 mutant mice

MiR-126^+/-^ mice were obtained from Dr. Eric Olson’s lab (Wang et al. 2008), were backcrossed into the C57BL/6 background (Sharma et al. 2019), and were maintained according to protocols approved by the Institutional Animal Care and Use Committee of Weill Cornell Medicine. The mice were housed in positive-pressure air-conditioned units (25°C, 50% relative humidity) on a 12 hour light:dark cycle and provided mouse chow and water *ad libitum*. MiR-126^-/-^ mice die between embryonic stage 15.5 (E15.5) and E18.5, due to vascular malformations and edema (Sharma et al. 2019). MiR-126^+/-^ males and females were intercrossed to yield E13.5 embryos that are wild-type, heterozygous, and homozygous for miR-126. Genotyping was performed by genomic PCR on DNA from yolk sac or tail biopsies as described previously (Sharma et al. 2019).

### Cell culture

iPSC-derived (New York Stem Cell Foundation iPSC library, NY) NPCs were kindly provided by Shuibing Chen (Weill Cornell Medicine). NPCs were cultured in 10 cm dishes, multi-well plates or on cover slips that were coated with growth factor reduced-Matrigel (Corning) (1:100 dilution in cold sterile PBS), in 50% Neurobasal medium and 50% Advanced DMEM/F12 (Gibco) supplemented with 1x neural induction supplement (Gibco). NPCs were differentiated into motor neurons in DMEM/F12 supplemented with 1x N2 and 1x B27 containing vitamin A (Gibco), 200 ng/µl fibroblast growth factor (Tonbo Bioscience), 200 ng/µl brain-derived neurotrophic factor (Peprotech), 200 ng/µl glial cell-derived neurotrophic factor (Peprotech), 200 ng/µl Neurotrophin-3 (Peprotech), 200 nM ascorbic acid (Rockville), 1 µg/ml laminin (Invitrogen) and 500 µg/ml N6,2’-O-Dibutyryladenosine 3’,5’-cyclic monophosphate sodium salt (dbcAMP, Sigma). Cells were used for experiments up to passage 8. The cells were maintained under standard cell culture conditions at 37°C in humidified air with 5% CO2. NPC identity was confirmed by examining cells for induction of Olig2 and Islet1 expression.

### Lentivirus-mediated overexpression and knock down of miR-126 in neural progenitor cells

HEK293T cells were co-transfected with lentivirus plasmid constructs for overexpression and knockdown of miR-126, and their respective controls, provided by Dr. John Dick (Lechman et al. 2012), together with packaging plasmid (pMDLg/pRRE), VSV fusogen plasmid (pVSV-G), and pRSV-Rev plasmid respectively, at a 4:2:1:1 ratio, using HEPES/CaCl2 precipitation. 48 and 72 hours after transfection, culture supernatants were collected and filtered to remove dead cells. Virus was concentrated 10-fold in neural expansion medium using Lenti-X concentrator (Takara) overnight at 4°C. NPC were infected at 80% confluency by adding virus stocks at a final dilution of 1:2 together with 4 µg/mL Polybrene (Santa Cruz), followed by spinning down of virus onto the NPC at 2300 rpm for 1 hour at 30°C. Neural expansion medium was replaced 4-6 hours after centrifugation. Transduction efficiency was determined by qRT-PCR and by immunofluorescence (IF) for RFP and GFP.

### qRT-PCR

Lentivirus-transduced NPCs were seeded into 6-well plates at 2.5×10^5^ cells/well in neural expansion medium and allowed to adhere overnight. NPCs were provided with fresh neural expansion medium or differentiation medium, cultured for 3 days and lysed in TRIzol (Invitrogen) for 5 minutes at room temperature. After chloroform (IBI Scientific) was added to the lysates for phase separation, samples were vortexed and centrifuged at 12,000 x g for 15 minutes at 4°C. The upper aqueous layer was transferred to a fresh tube, mixed with 100% isopropanol and centrifuged at 12,000 x g for 10 minutes at 4°C. Pellets were washed twice in ice-cold 70% ETtOH and centrifuged at 12,000 x g for 5 minutes at 4°C. RNA pellets were air-dried for 15 minutes and resuspended in RNAse-free water, treated with DNAse (Ambion) for 15 minutes at 37°C, followed by DNAse inactivation (5 minutes at 65°C). Purified RNA was reverse transcribed using qScript (QuantaBio) and qRT-PCR was performed using PerfeCTa SYBR Green Supermix for iQ (QuantaBio) and primers (Table 1) in a iQ5 iCycler (BioRad). For detection of miR-126 and the control mRNA sno234, purified RNA was reverse transcribed and qRT-PCR was performed using TaqMan MicroRNA Assays (#001234 for miR-126 and #000450 for sno234, Applied Biosystems) and Universal PCR Master Mix no AmpErase UNG (Applied Biosystems).

**Table 1:**
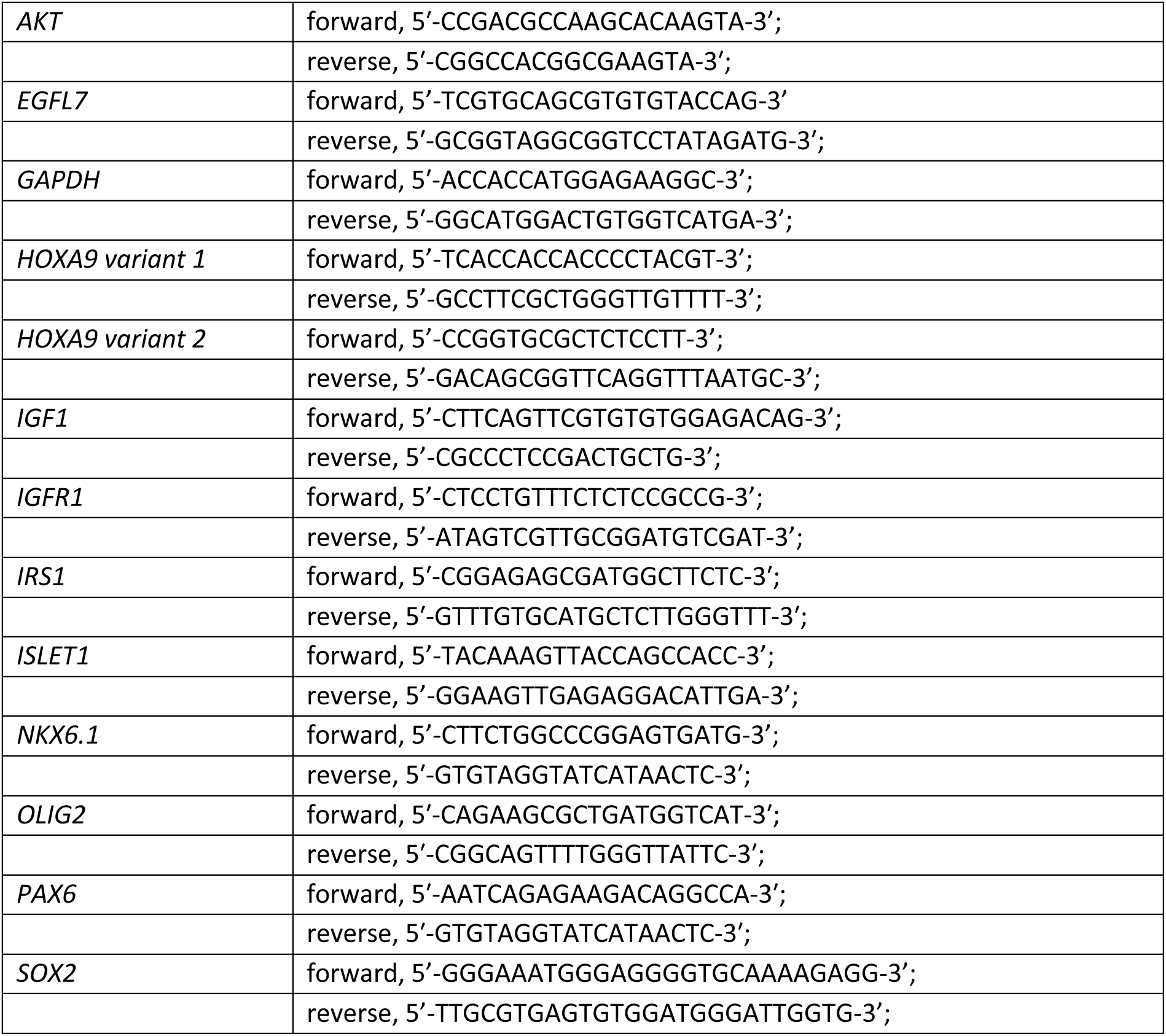
Primers.

### Immunofluorescence staining

For timed pregnancies, the day of the vaginal plug was counted as embryonic day 0.5 (E0.5). E13.5 embryos were dissected from the uterus of the pregnant female, fixed in 4% paraformaldehyde, embedded in OCT and cryosectioned as described (Sharma et al. 2019). Slides containing cryosections were rinsed with PBS, followed by blocking with PBST containing 10% donkey serum for 1 hour at room temperature. Sections were subjected to immunofluorescence staining by incubating with primary antibodies overnight at 4°C, followed by washing with cold PBS (three times for 3 minutes each) and incubation with secondary antibodies in PBST containing 10% donkey serum for 2 hours at room temperature. Primary and secondary antibodies are listed in Tables 2 and 3. After washing twice with cold PBST and twice with cold PBS, nuclei were stained using Prolong Gold antifade reagent with DAPI (Invitrogen) or HCS NuclearMask Deep Red Staining (Invitrogen). Images were obtained using an AxioPlan2 (Zeiss) microscope, and data were analyzed using ImageJ. For immunofluorescence staining of NPCs, NPCs were seeded at 5×10^4^ cells on coated cover slips in 24-well plates in fresh neural expansion medium and allowed to adhere overnight. NPCs were rinsed with ice-cold PBS and fixed with 4% paraformaldehyde for 10 minutes at room temperature, followed by blocking with PBST containing 10% donkey serum for 1 hour at room temperature. Immunofluorescent staining was done as described above.

**Table 2:**
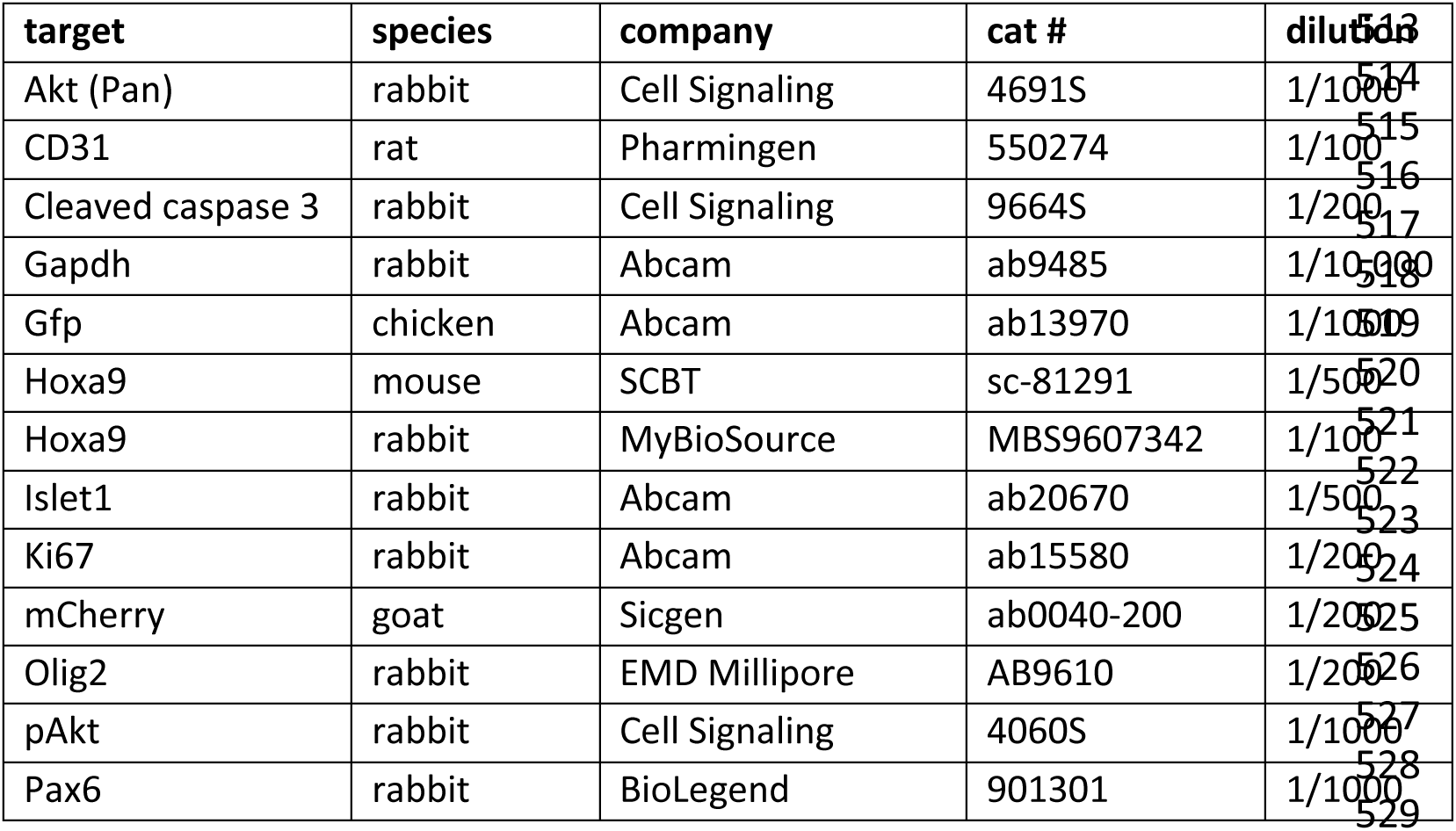
Primary antibodies.

**Table 3:** Secondary antibodies.

| Host/target | label | company | dilution |
| --- | --- | --- | --- |
| Donkey anti-chicken | Alexa488 | Jackson labs | 1/200 |
| Donkey anti-goat | Alexa488 | Jackson labs | 1/200 |
| Donkey anti-rabbit | Alexa488 | Jackson labs | 1/200 |
| Donkey anti-chicken | Alexa555 | Abcam | 1/200 |
| Donkey anti-rabbit | Alexa555 | Abcam | 1/200 |
| Donkey anti-rat | Alexa555 | Abcam | 1/200 |
| Donkey anti-mouse | HRP | Jackson labs | 1/10,000 |
| Donkey anti-rabbit | HRP | Jackson labs | 1/10,000 |

### Western Blotting

1×10^6^ lentivirus-transduced NPCs were plated into 10 cm dishes containing neural expansion medium and allowed to grow to 90% confluency. Cells were lysed in ice-cold RIPA buffer supplemented with 1x complete protease inhibitor cocktail (Roche) and PhosSTOP phosphatase inhibitor (Roche), and protein concentration of the lysates was determined using BCA Protein Assay Reagent (Pierce). 30-40 micrograms of proteins were separated on 12% SDS-PAGE gels and transferred to polyvinylidene fluoride (PVDF) Immobilon-P transfer membranes (Millipore). After washing in PBS and blocking in blocking buffer (5% BSA in PBST) for 1 hour at room temperature, membranes were incubated with primary antibodies in blocking buffer overnight at 4°C. Membranes were washed 3 times with PBS, incubated with secondary antibodies in blocking buffer (2.5% BSA in PBST) for 1 hour at room temperature, washed 3 times with PBS and exposed to film (ThermoFisher) using a horseradish peroxidase (HRP) substrate for enhanced chemiluminescence (ECL) (Pierce). Intensities of bands were analyzed using Image J. Primary and secondary antibodies are listed in Tables 2 and 3.

### ELISA

miR-126 transduced NPCs were seeded 1×10^6^ cells/well into six-well plates in fresh neural expansion medium and cells were allowed to grow ∼90% confluent. Culture supernatants were treated with 0.25N HCl to lower the pH to 3 in order to dissociate IGFBP-bound IGF-1. Recombinant human IGF-2 (100 µg/ml) was added to compete with IGF-1 for re-binding to the free IGFBPs, after adding 1N NaOH to return the pH to 8 (Baxter 2001). pH-indicator strips (ColorPhast) were used for determining the pH. Treated supernatants were analyzed for human IGF-1 by ELISA according to manufacturer’s instructions (RayBio, Norcross, USA).

### Statistical analysis

Data were analyzed using the Kruskal-Wallis (One-way ANOVA) test followed by the Mann-Whitney U test. A *P*-value < 0.05 was considered significant.

